# GPS-Net: discovering prognostic pathway modules based on network regularized kernel learning

**DOI:** 10.1101/2024.07.15.603645

**Authors:** Sijie Yao, Kaiqiao Li, Tingyi Li, Xiaoqing Yu, Pei Fen Kuan, Xuefeng Wang

## Abstract

The search for prognostic biomarkers capable of predicting patient outcomes, by analyzing gene expression in tissue samples and other molecular profiles, remains largely on single-gene-based or global-gene-search approaches. Gene-centric approaches, while foundational, fail to capture the higher-order dependencies that reflect the activities of co-regulated processes, pathway alterations, and regulatory networks, all of which are crucial in determining the patient outcomes in complex diseases like cancer. Here, we introduce GPS-Net, a computational framework that fills the gap in efficiently identifying prognostic modules by incorporating the holistic pathway structures and the network of gene interactions. By innovatively incorporating advanced multiple kernel learning techniques and network-based regularization, the proposed method not only enhances the accuracy of biomarker and pathway identification but also significantly reduces computational complexity, as demonstrated by extensive simulation studies. Applying GPS-Net, we identified key pathways that are predictive of patient outcomes in a cancer immunotherapy study. Overall, our approach provides a novel framework that renders genome-wide pathway-level prognostic analysis both feasible and scalable, synergizing both mechanism-driven and data-driven for precision genomics.

## Introduction

In biomedical research, the advent of high-throughput technologies has introduced an abundance of high-dimensional data, especially in genomics. However, the analysis of such high-dimensional biological data poses several challenges. Traditional statistical and machine learning methods, like Lasso ^1^ or Elastic Net ^2^, which are widely used for biomarker selection, often focus on individual genes or features. This gene-level approach, while valuable, overlooks the broader context of biological pathways and networks. It does not fully capture the interactions and dependencies among genes and proteins, which are essential for understanding the functioning of complex biological systems.

In response to these limitations, a network-constrained regularization model that enriches biomarker selection was proposed ^3^. This model introduces graphical structures into the framework. The core of this network regularization utilizes the Laplacian matrix, a fundamental element that describes the connections and relationships within biological networks. The Laplacian matrix, derived from the adjacency matrix of the network, captures how genes interact and coordinate their activities within biological pathways. Although this model does incorporate biological pathway information, its primary focus remains on selecting markers at the gene level. In this context, it is important to acknowledge the significance of biological pathways, often referred to as gene sets, which represent collections of genes jointly engaged in specific biological processes. These pathways are fundamental components of the complex biological network, and they play a pivotal role for coordinating biological functions. Consequently, the ability of network regularization models to discern significant pathways in cases where multiple pathways are present in the data is limited.

In genomics data analysis, the organization of genes or variables into functionally related or co-expressed groups, such as pathways, is a common practice. The Molecular Signatures Database (MSigDB) ^4^ stands out as one of the largest and most widely used repositories of gene sets, including nine human collections denoted as C1-C8 and hallmark gene sets. To address the natural group structures in genomics data, the Group Lasso ^5^ was proposed to extend the Lasso to group level, allowing entire groups of correlated variables to be selected or excluded together. The number of pathways or gene sets differs among these collections. The hallmark collection, with 50 gene sets, can be efficiently managed by Group Lasso for pathway selection. Conversely, collections like ontology gene sets (C5) and immunologic signature gene sets (C7) contain an excess of 5000 gene sets (C5: 15937 sets and C7: 5219 sets). In such instances, the efficiency of Group Lasso experiences a notable decline due to the need for regularization in each group during the optimization process. Although Group Lasso can be employed for pathway selection, its limitation lies in the absence of integration with biological information. Additionally, it cannot be directly applied to pathways with gene overlaps, which frequently happen in biological pathways. These limitations restrict its capability to effectively select genome-wide pathways.

To address the computational challenges posed by the large number of pathways, there has been a shift in focus toward leveraging the flexibility offered by Multiple Kernel Learning (MKL) ^6,7^. MKL is an extension of support vector machines (SVMs) ^8^ that enables the combination of information from multiple kernels. In contrast to SVMs, which elevate the original data dimension to find a hyperplane for optimal data point separation in the feature space, our emphasis lies in the capacity of MKL to reduce feature dimensions in pathway selection ^9-11^. MKL proves to be efficient in integrating information from diverse sources or representations in genomic data ^12^. Particularly, the pathway selection process involves transforming different pathways into distinct kernels, which is achieved through the application of various kernel functions. Notably, there are three commonly employed kernel functions: the linear kernel, Gaussian kernel, and graph kernel. The linear kernel, while straightforward to interpret, possesses limitations in its capacity to capture non-linear relationships within the data. Conversely, the Gaussian kernel excels in capturing both linear and non-linear relationships, providing a versatile approach. However, it lacks the capability to incorporate biological knowledge into the modeling process, resulting in less interpretable prediction models. In contrast, the graph kernel approach stands out by integrating biological information. In this study, we define a set of pathway-network graph kernel functions by incorporating the Laplacian matrices. This approach allows each kernel to include the prior knowledge, thereby enhancing the potentials in large number pathway selection. This characteristic aligns seamlessly with the network regularization model, in which the kernel can be constructed using network information (the Laplacian matrix) in conjunction with data. Additionally, the computational efficiency of employing MKL for the selection of numerous pathways can be further enhanced by leveraging specialized algorithms like SpicyMKL which utilizes the proximal minimization on a sparse kernel combination ^13^.

To employ the MKL techniques in pathway selection in cancer genomics data, extension for including multiple pathways is needed based on the previously mentioned network regularization model. The bi-network regularization model ^14^ was originally proposed for the scenario that genes within the dataset could manifest two distinct network structures. For instance, one structure may arise from the inherent data characteristics, reflecting a data-driven network structure. Conversely, the other structure could mirror well-established biological network configurations. In the real world, it is often unrealistic to pinpoint the actual network structure, hence a reasonable assumption is that the real network can be approximated by combining these two distinct structures. The fusion of two diverse structures not only harnesses the information embedded in the dataset but also serves as a vital link to the prior biological knowledge. In addition, the bi-network regularization model can also be applied when the data contains two different biological pathways, such as two distinct gene sets. Extending from this application of bi-network regularization model for two pathways, it is natural to consider the expansion from utilizing two networks to multiple networks for pathway selection.

In this study, we introduce the Genome-wide Pathway Selection with Network Regularization (GPS-Net) which extends bi-network regularization model to multiple-network and employs multiple kernel learning (MKL) for pathway selection. In pathway selection, GPS-Net deploys the pathway-kernel (PK) approach to efficiently manage numerous pathways. By converting each pathway into a distinct kernel using graph kernel functions, PK streamlines pathway selection through the efficient MKL algorithm, eliminating the need for exhaustive computations. GPS-Net harnesses the logistic model for binary responses and the Cox model for survival outcomes to navigate the pathway and gene selection process. To improve gene biomarker selection accuracy, GPS-Net employs the augmented penalized minimization of *L*_0_ (APM-*L*_0_) approach ^15^. This added *L*_0_ penalization substantially reduces false positives, particularly in scenarios with a limited number of significant biomarkers. While the use of MKL in the logistic model has been widely explored, its application in the Cox model remains a relatively underexplored area. In this study, we focus on the application of GPS-Net in the analysis of survival data, offering new insights into the exploration of network regularization and the MKL approach within the Cox model. Our research serves not only to highlight the practicality and robustness of GPS-Net but also to emphasize its effectiveness in selecting significant pathways and biomarkers. We demonstrate these advantages through simulated experiments and real-data analyses, highlighting the power and versatility of GPS-Net in pathway selection within the complex high-dimensional genomics data.

## Material and methods

### Overview of GPS-Net

We introduce GPS-Net, a user-friendly framework designed for the selection of pathways within high-dimensional genomics data. GPS-Net operates by taking genomics data X, along with its response Y, and incorporates pathway information as input. The pathway information is a gene set collection which can be sourced from MsigDB or manually constructed from users’ preferences. To integrate pathway information into the model, GPS-Net initially extracts the network structure for each pathway, typically represented by Laplacian matrix derived either through data-driven computation or sourced from established biological pathway databases like STRING^16^. We offer a function in GPS-Net package that utilizes the STRING API to retrieve the STRING gene interaction network. Subsequently, GPS-Net reconstructs each network kernel with one Laplacian matrix, thereby transforming the pathway selection problem into a multiple kernel learning (MKL) process. By solving the MKL problem, GPS-Net identifies and selects kernels corresponding to specific pathways. Finally, GPS-Net converts the MKL estimators back to the original pathway selection coefficients, resulting in an output that contains the selected pathways and the estimated coefficient vector 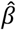. The workflow of GPS-Net is depicted in Figure 1.

**Figure 1.**
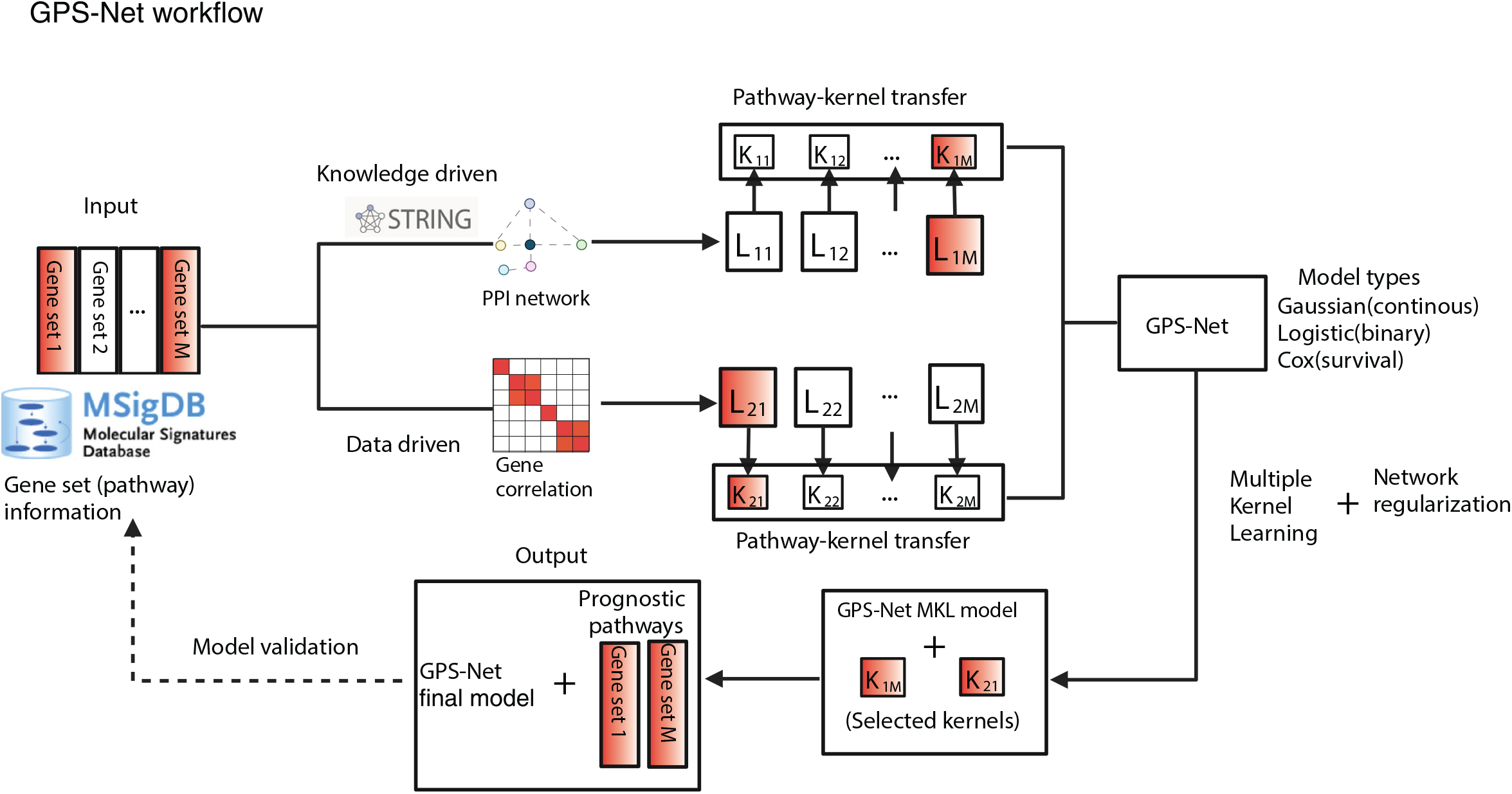
Overview workflow of GPS-Net.

### Network regularization model

To provide a foundational understanding of the genome-wide pathway selection with network regularization in GPS-Net, we begin by considering a linear lasso regression model ^1^ with n-dimensional response Y, predictors (genes) *X* = (*X*_1_, …, *X*_*P*_) ∈ ℝ^*n*×*p*^. The objective function is defined by adding an *L*_1_ penalty to the conventional linear regression model:

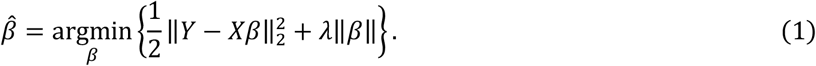

The Lasso estimator offers a sparse solution, effectively functioning as a variable selection mechanism. Transitioning from the linear model (1) to the Generalized Linear Model (GLM) is straightforward, where we replace the squared error term 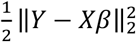 in Equation (1) with the negative log-likelihood of the model, denoted as (*β*). In this study, we specifically focus on survival data, which is routinely examined in cancer research for biomarker discovery. Therefore, *ℓ*(*β*) can be considered as the partial log-likelihood of the Cox model ^17^. To improve the model performance, various selection penalties are introduced such as the Elastic Net ^2^, the MCP ^18^, and the SCAD ^19,20^. However, these methods tend to overlook the estimation of specific information pertaining to correlated patterns among genes *X*, which has the potential to enhance the accuracy of estimations and predictions. To connect with the pairwise structure among genes represented by *X*, a network-constrained and graph-constrained estimation model was proposed by combing the *L*_1_ penalty and a Laplacian quadratic penalty ^3^:

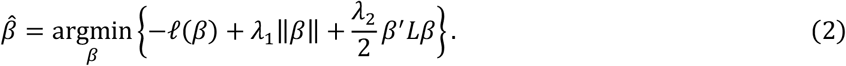

Here *L* is a normalized Laplacian matrix. The biological pathway or network is depicted as a graph, with genes serving as nodes and their interactions as edges. The Laplacian matrix derived from this graph reflects the pairwise associations between genes within the network. The matrix is constructed in a way such that genes connected by an edge have non-zero entries, whereas unconnected genes have zero entries. Details of the mathematical definition of normalized Laplacian matrix L is given in **Supplemental methods A.1**. Notably, if L is an identity matrix, the problem (2) transforms into an adaptive elastic net problem. It’s worth mentioning that the Laplacian matrix L can be constructed from data or predetermined using existing knowledge. Several prior studies have examined the methods and solutions for graph-constrained regularization problems (2) ^21, 22^. However, these studies have primarily focused on the application of a single Laplacian matrix penalty. In the following section, we will introduce a novel model that considers the cases involving two Laplacian matrices penalty, corresponding to instances with two distinct network structures.

### Bi-network regularization model

In practical data analysis, acquiring the precise network structure as represented by the Laplacian matrix poses significant challenges and often proves unattainable. Prior studies primarily focused on employing Laplacian matrices derived solely from the data itself ^22^. In this study, the network directly computed based on the correlation structure of the input gene expression data is referred to as reference-free network. We then introduce referenced-based net to incorporate prior biological knowledge into the model. This network refers to a biological network derived from established gene relationship databases such as STRING^16^, KEGG ^23^ and Reactome ^24^. To integrate both reference-based and reference-free networks, we introduce below a bi-network regularization model ^14^:

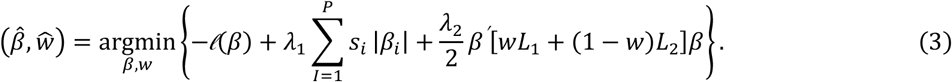

Here *s* = (*s*_1_, …, *s*_*p*_) is the weight vector of L-1 penalty, defined similarly to the adaptive lasso framework. The weights *s*_*i*_ can be pre-estimated by 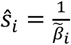 where 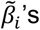 are the estimators by solving (3) with *λ*_2_ = 0 25,26 The final network representation, denoted as *L*, is obtained as the weighted sum of both *L*_1_ and *L*_2_. The weight *w* is assigned to incorporate gene interaction and correlation information from both the reference-based and reference-free networks, as defined previously. To ensure the positive semi-definiteness of the Laplacian matrix *L*, the value of *w* ranges from 0 to 1. This integrated network builds upon a similar idea proposed in the Net-Cox model ^27^, which utilizes a weighted sum of the Laplacian matrix and the identity matrix. The bi-network regularization model allows the integrated network model to incorporate either the data-driven (reference-free) or existing database-driven (reference-based) network (when *w* = 0 or *w* = 1), or a hybrid (0 < *w* < 1) network. This hybrid approach more accurately captures the relationships among all genes in the pathway or set, leading to improved performance in pathway selection and thus final prediction accuracy.

To solve the problem (3), there is an additional parameter *w* to estimate. However, the conventional way for solving optimization problem by taking the first derivatives of the objective function to zero is inapplicable when it comes to estimating *w*. To tackle this problem, a cross-validation (CV) method was recommended by treating *w* as an extra tunning parameter similar to *λ*_1_, *λ*_2_. The details of the solution for problem (3) are provided in the **Supplemental methods A.2**.

We introduce bi-network regularization model to improve the efficacy of biomarker selection by combining prior biological knowledge with data driven information. However, since model (3) shifts from employing a single Laplacian matrix in network regularization models to two, it naturally allows for expansion to include more than two networks, thereby enabling the model to integrate information from multiple pathways. This is achieved by replacing the summation of two Laplacian matrix penalties with a composite of multiple Laplacian matrices 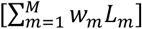, where each Laplacian matrix encapsulates distinct gene network information from different pathways. In this scenario, the weight parameter *w* becomes an M-dimension vector and the searching space over (*w, λ*_2_) expands exponentially. Although the CV method can handle cases with limited M (e.g., M=3 or 4) to estimate a final integrated Laplacian matrix, the pathway selection process in estimating *w* becomes computationally burdensome. This computational burden persists even when dealing with the hallmark collection, which is the smallest among MSigDB human gene sets collections, with M=50 gene sets (pathways). To address this issue, we proposed the genome-wide pathway selection with network regularization (GPS-Net) framework in the following section.

### Genome-wide pathway selection with network regularization framework

#### Multiple pathways

Expanding on the knowledge and experience learned from the bi-network regularization model, we further investigated the pathway selection problem in cases involving multiple pathways. In real-world scenarios, pathway information can consist of more than two candidate structures 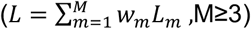. This, however, presents a new challenge as the dimension of the weight searching space increases exponentially if the CV procedure used in (3) is employed. We emphasized the pathway selection by introducing a network elastic net model for multiple pathways:

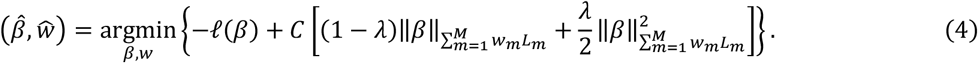

To ensure the semi-positive definiteness of the combined Laplacian matrix 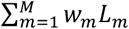, the weights *w*_*m*_ should be non-negative, and are subject to the constraint that they sum to one. In equation (4), a composite penalty, comprising a non-smooth grouped penalty 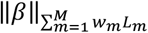 and a quadratic penalty 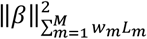, has been devised to enhance the accuracy of pathway selection. Here, 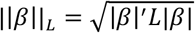 and 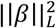 is the quadratic of 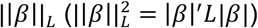. In the multiple pathway selection model, the focus shifts from the individual gene level to the pathway level, thus only group-wise penalties are applied in (4). Genes within the same pathway are likely to share biological functions. Therefore, incorporating the group selection model can enhance selection accuracy where genes exhibiting similar behavior within the same pathway are selected together. Notably, the problem (4) can be reduced to a group lasso ^5^ framework by setting all *L*_*m*_^′^*s* as identity matrices and *λ* = 0, eliminating the network structure and the quadratic penalty. Additionally, problem (4) also confronts the challenge of finding the optimal *w*_*m*_ with CV procedure. The computational cost increases exponentially with the number of pathways M. To tackle this challenge, we introduced the pathway-kernel approach to solve (4).

#### Pathway-kernel approach

To efficiently handle pathway selection with multiple candidate structures, a pathway-kernel (PK) approach is introduced. In this approach, each pathway (*L*_*m*_) is transformed into its own kernel using specific graph kernel functions 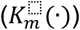. This generates kernels 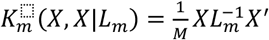 for each pathway.

The dimension of *K*_*m*_ is *n* × *n*, and by applying 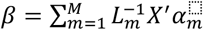, problem (4) becomes a multiple kernel learning (MKL) optimization problem:

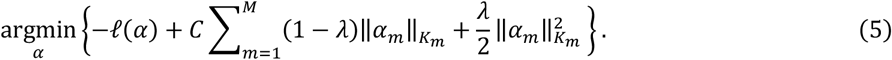

The choice of kernel function transforms the original p-dimensional optimization problem to an equivalent n-dimensional dual optimization problem, thereby enhancing computational efficiency. Additionally, each kernel is constructed by normalizing the original feature space X with 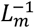. Hence, we can solve the MKL problem (5) to directly select the functional pathways. The advantage of the PK approach is that it simplifies the gene-pathway selection issue by making pathway selection equivalent to the sparsity of kernel coefficient *α*. The regularization procedure in MKL optimization forces sparsity in the estimated kernel coefficient 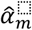, leading to the selection of functional gene biomarkers and pathways. The corresponding pathway weights 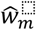 can then be calculated from estimated kernel coefficients. The solution for the PK approach used in (4) includes the dual optimization in MKL process. For detailed steps, refer to **Supplemental methods A.3**.

To further improve the pathway selection performance in the PK approach, we apply an adapted version of the augmented penalized minimization of *L*_0_ (APM-*L*_0_) ^15^ step in (5). This adjustment effectively reduces the occurrence of false positives, leading to more accurate pathway selections. The adapted APM-*L*_0_ with SpicyMKL is given in **Supplemental methods A.4**.

#### GPS-Net implementation

To leverage the models described above, we introduced the Genome-wide pathway selection with network regularization (GPS-Net) framework. GPS-Net framework integrates the multi-pathway-PK procedure for pathway selection with the bi-network regularization model, harnessing both empirical data and prior biological knowledge.

Within the GPS-Net framework, the initial step involves parallel pathway selection using reference-free and reference-based networks, employing the PK approach as described in equation (4) and (5). The reference-free Laplacian matrices are calculated through the partitioning of pathways in data X and its correlation matrix. Meanwhile, the reference-based networks can be sourced from external biological network references like STRING, KEGG, and Reactome. Upon the completion of pathway selection, two integrated Laplacian matrices are derived : 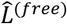 and 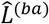, where 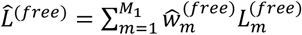 and 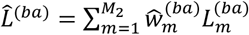.

The final step in the process is to employ these integrated Laplacian matrices 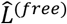 and 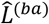 within the bi-network regularization model, introducing a balance parameter denoted as *γ*,

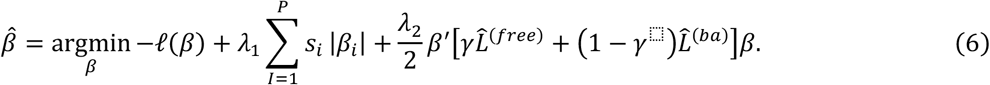

The value of *γ* determines the balance between data-driven and prior-knowledge based networks. The default value of *γ* is 0.5, indicating no preference between data-driven and prior-knowledge based networks. The CV procedure introduced for the bi-network regularization model (3) can also be applied in this step to find a computational optimal *γ*.

A key feature of GPS-Net is its reliance on pathway information or group partition {*P*_1_, …, *P*_*M*_|*P*_*m*_ ⊂ *X*} as input, providing researchers the flexibility to either calculate reference-free pathway network structures or input their own pre-assigned networks or even a combination of both. The implementation workflow of reference-based GPS-Net is detailed in **Supplemental methods A.6**.

## Results

### Simulation scenarios

To comprehensively evaluate the effectiveness of the GPS-Net framework, we conducted a series of simulation studies. These investigations aimed to compare GPS-Net with several existing biomarker selection methods, including Elastic-net (Enet), Lasso, APM-*L*_0_ Net (ANet) and Group Lasso (GLasso). The objective was to gain insights into GPS-Net’s performance across diverse scenarios and dataset types, allowing for a thorough assessment of its strengths and potential areas for improvement.

We simulated 500 samples in each scenario, while considering different combinations of feature dimensions (P) and pathway numbers (M). In addition, three diverse network structures which reflect different levels of correlations among genes within pathways were investigated. The covariate data X was generated leveraging the network structures assigned to the pathways using R package “huge” ^28^. Specifically, we configured the off-diagonal elements of the covariance matrix to values of 0, 0.3, and 0.8 corresponding to three network structures: independent, medium-correlated and highly-correlated. Survival data was generated using the Cox model, with 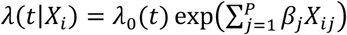. Each simulation scenario was repeated 100 times, and an approximate 30% censoring rate was applied. The values of these non-zero *β* coefficients were defined as 2 × (−1)^*q*^, where q followed a Bernoulli distribution with a probability of 0.5.

1. In Scenario1, we simulated a small set of pathways (M=50), matching the size of hallmark gene sets. Each pathway consisted of 200 genes, resembling the structure of hallmark gene sets, which has a median gene number of 180. No overlapping genes were allowed across the 50 pathways. We designated 3 pathways as significant, and within each significant pathway, 5 β values were set as non-zero. We assessed the methods’ performance using metrics including the out-of-sample concordance index (C-index), group accuracy, specificity, sensitivity, and precision.
2. In Scenario 2, we simulated an intermediate number of pathways (M=500), close to the size of C1 (301). The median number of genes in C1 gene sets is 115, so we generated each pathway with 100 genes, resulting in a total of 50,000 genes without overlap. Among the 500 pathways, we designated 5 as significant, each with 10 non-zero β values out of 100. To evaluate Scenario 2, we calculated the C-index, sensitivity, and precision similarly as in Scenario 1. We further investigated the impact of different Laplacian matrices used in GPS-Net on pathway selection. In this simulation, we only considered the reference-free network case since we were not generating real biological pathways. We evaluated GPS-Net in three different contexts: 1. True Laplacian matrix (GPS-T); 2. estimated reference-free Laplacia matrix (GPS-ES); 3. Identity matrix (GLasso). The estimated reference-free Laplacian matrices are calculated by estimating the precision matrix which is the inverse of the covariance matrix with ‘get_free_net’ function in GPS-Net package. The procedure for estimating the reference-free Laplacian matrices with precision matrix is given in **Supplemental methods A.5**. Notably, as we applied different algorithms and an extra *L*_0_ regularization in GPS-Net, the performances between GPS-T and Glasso in independent scenario are not identical.
3. 3. When the number of pathways increases, it is natural to have overlap genes in different pathways. In Scenario 3, we start to consider the overlapping genes with large number of pathways. We simulated the 1000 pathways, similar to the size of C8 (829), with each pathway containing 100 genes. We set the total number of genes to 20,000, comparable to the 20,481 genes in C8. Each pathway is assigned 20 unique genes and 80 random overlaping genes. The first 5 pathways are designated as significant with each containing 10 non-zero β.
4. 4. In Scenario 4, we aimed to validate GPS-Net’s pathway selection capabilities in a real-world application. This scenario included a significant hypoxia ^29^ alongside a set of another M =5,10 or 50 synthetic pathways as background. 100 replicates were conducted to calculate the frequency of hypoxia pathway selection. It has been shown that tumors in the head and neck region can experience hypoxic conditions, leading to the activation of hypoxia-inducible factor (HIF) ^30-32^. We hypothesis that the GPS-Net can successfully identify the hypoxia pathway among a collection of background pathways.
5. 5. In Scenario 5, we conducted a type I error analysis with an ultra-high number of pathways (M=10,000) to reflect the structure of GO gene sets, which consist of 10,460 sets. For each selection method (GPS-NET, ANet, ENet, Lasso, GLasso), the type I error was calculated with a permutation test across different number of pathway settings ^33^. Specifically, we generate the data with two different models: (1) a null model where all *β* = 0, and (2) a mixed model with the first 5 pathways each containing 5 non-zero *β*=2 × (−1)^*q*^. All data were generated by setting the off-diagonal element of sparse correlation matrix to 0.5. The simulation pathway settings were summarized in **Table S1**.

### GPS-Net’s performance across simulations

In Scenario 1, GPS-Net and ANet emerged as the top performers, consistently achieving average C-index values above 0.8. GPS-Net slightly outperformed ANet due to the additional pathway information considered, achieving the highest average C-index value of 0.88, which is closely followed by ANet with 0.85. Other methods, including Enet, Lasso, and GLasso, all had C-index values below 0.75. As the complexity of network structure increases, we observed a decline in performance across all methods. Enet and Lasso performed similarly, with C-index values ranging from 0.75 to 0.7, while Glasso performed the worst in this scenario. Notably, GPS-Net showed the most robust performance with the least decline in C-index values (Figure 2A). We also compared the average performance across three network structures, using metrics including group accuracy, specificity, sensitivity, and precision. Sensitivity, and specificity were all close to 1 for all the methods, indicating that true positives can be mostly selected by any methods in Scenario 1. GPS-Net and ANet maintained high precision values above 0.9, whereas other methods suffered from high false positives, resulting in precision values lower than 0.25 (Figure 2B). Only GPS-Net and GLasso possessed the functionality for group selection, with GPS-Net outperforming GLasso (0.92 to 0.85) in Scenario 1 when the pathway number is not large.

In the simulation Scenario 2, GPS-Net consistently achieved the highest C-index values, measuring 0.69, 0.66, and 0.65, respectively. ANet remained the second-best with C-index values of 0.63, 0.61, and 0.60. In contrast, both Lasso and Enet returned average C-index values below 0.6. Notably, Glasso exhibited relatively improved performance in Scenario 2 compared to Scenario 1, surpassing Lasso and Enet with a C-index value of approximately 0.6 (Figure 2A). Compared to Scenario 1, Scenario 2 showed a decreased C-index across all methods, with GPS-Net declining the least. This trend highlights the stability of GPS-Net in delivering consistent performance for the case with an intermediate number of pathways (Figure 2A). In addition, the superior performances of GPS-Net was evident in the average selection accuracy observed in Scenario 2 (Figure 2C). GPS-Net showed a relatively high precision of 0.78 comparing to Enet (0.46), Lasso (0.42) and Glasso (0.53). Both GPS-Net and Glasso were efficient in selecting most true positives, with sensitivity values above 0.7. In terms of pathway selection performances, three methods (ANet, Enet, and Lasso) that lack pathway selection capability, showed accuracy rated at 0 as expected. GPS-Net outperformed Glasso in Scenario 2 (0.75 to 0.69), indicating that GPS-Net successfully identified over 70% of the true pathways, whereas Glasso identified less than 70% of the true pathways.

We further tested the robustness of GPS-Net using estimated Laplacian matrix as the regularization term. In the independent case where the true network is represented by the identity matrix, GPS-T and GLasso use the same original pathway selection model but employ different algorithms. GPS-T and GLasso performed similarly to each other in this case, with higher F1 scores (>0.8) compared to GPS-ES (0.76). The high F1 scores of GPS-T across all three categories suggests that employing the correct network regularization significantly enhances the precision of pathway selection. However, as the complexity of the network structure increased, GPS-ES’s performance became competitive with GPS-T. In scenarios with medium correlation, GPS-ES outperformed GLasso with an F1 score of 0.75 compared to GLasso’s 0.71. In highly correlated scenarios, GPS-ES maintained a F1 score of 0.74 compared to GLasso’s decreased F1 score of 0.68, making GPS-ES the second highest after GPS-T (Figure 2C). The selection performances are also evaluated using precision and sensitivity in different Laplacian matrix contexts (Figure 2C). GPS-T and GPS-ES demonstrated significantly higher precision compared to GLasso, exceeding 75% in both highly-correlated and medium-correlated network structures. In contrast, GLasso achieved only 65.6% for highly-correlated and 69.5% for medium-correlated network structures (Figure 2C). All methods showed similar sensitivity, due to the large number of true negatives simulated in this scenario. This trend underscores the adaptive nature of GPS-ES in coping with more complicated network structures, showcasing its robustness and effectiveness in real application where the true network is unknown.

**Figure 2.**
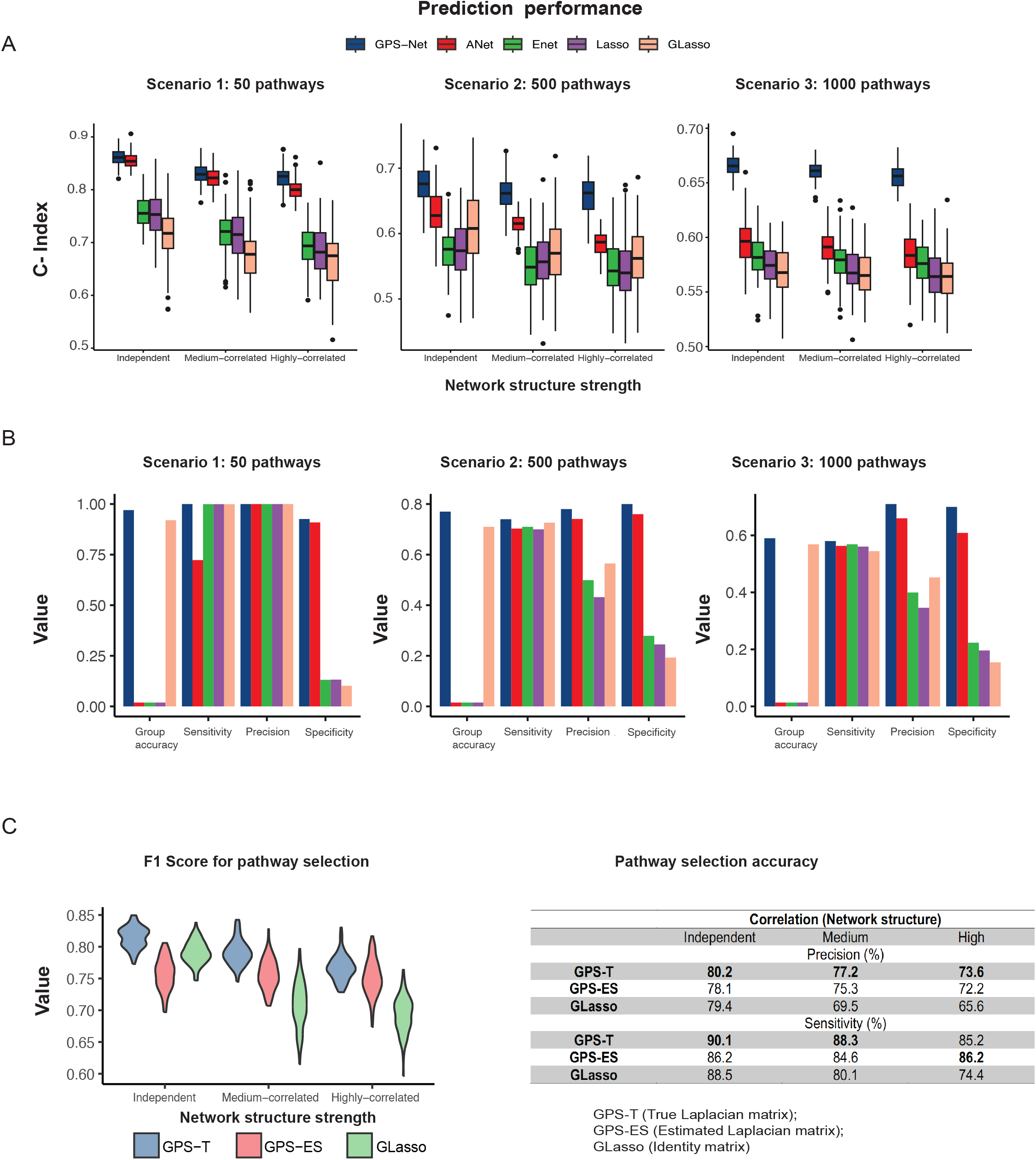
GPS-Net outperforms the other state-of-the-art selection methods in different simulation scenarios. (A) Predicted C-index from GPS-Net and other methods with increasing complexity of pathway structures for simulation Scenario 1 (left), Scenario 2 (middle) and Scenario 3 (right). (B) Average performances of GPS-Net and other methods in five selection measurements for Scenario 1 (left), Scenario 2 (middle) and Scenario 3 (right) (C) Pathway selection investigation of GPS-Net: F1 score measurement (left), Precision and Sensitivity measurement (right).

In Scenario 4, we simulated 100 replicates to evaluate GPS-Net in real data application with a special focus on the hypoxia pathway. The details of design scenario are given in Figure 3A. Within these experiments, the hypoxia pathway consistently emerges as a top choice in GPS-Net’s selections, being chosen 100, 78, and 54 times out of 100 with an increasing number of pathways (**Figure S1** and Figure 3B). In fact, it is the most frequently selected pathway when using GPS-Net. This finding demonstrates the robustness and reliability of GPS-Net in identifying pathways of biological significance with extra noisy information involved. However, in the Glasso selection results, when pathway number increases to 50, hypoxia is not the pathway most frequently chosen with 28 times out of 100. The remain 50 simulated pathways shows a more random pattern in their selection frequencies, with occurrences around 10 to 30 (Figure 3B). This observation highlights a notable disparity in the ability of GPS-Net and GLasso to capture pathways with biological significance, further underscoring the advantages of GPS-Net’s integration of additional biological network information.

**Figure 3.**
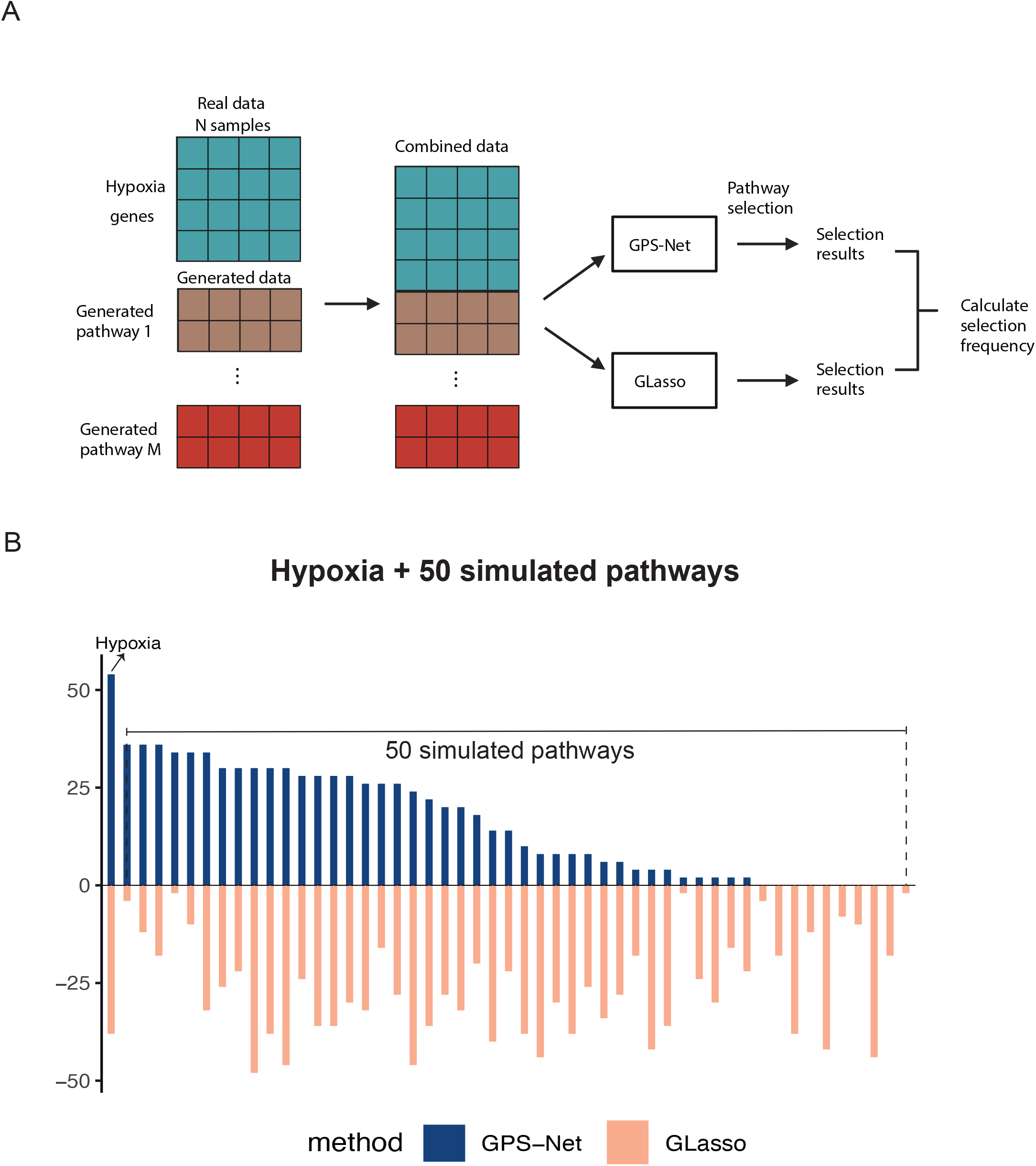
Targeted pathway (Hypoxia) selection comparison between GPS-Net and GLasso. (A) Experiment details of simulation Scenario 4. (B) Selection frequencies for hypoxia and other 50 generated pathways using real patients’ survival responses.

In Scenario 5, we calculated the type I error rates at significance levels of 0.001, 0.01, and 0.05, respectively. To reflect the structure of GO gene sets, we simulated M=10,000 pathways. All five methods maintained a low error rate in the null model as expected. However, in the mixed model (with non-zero *β*), a general increase in type I error rates was observed across all methods. Specifically, because the total gene number remained the same for both pathway number M=1,000 and M=10,000, the performances of single-gene selection methods (ANet, ENet, and Lasso) were very similar. In contrast, the group-wise methods (GPS-Net and GLasso) exhibited increased error rates, with GPS-Net from 0.165 to 0.184 and GLasso from 0.0253 to 0.0312 at a significance level of 0.001 (**Table S2**).

Together, these simulation scenarios represent a comprehensive assessment of the GPS-Net framework’s performance in various situations. They show GPS-Net’s efficacy in biomarker and pathway selection when compared to existing methods. The outcomes highlight GPS-Net’s proficiency in detecting significant pathways, higlighting its value for researchers in biomarker discovery and pathway selection. This significance is particularly evident in situations featuring complicated network structures of pathways.

### Validation of GPS-Net’s pathway selection proficiency with bladder cancer data and pathway references

The real-world application of the GPS-Net framework was conducted using the IMVIGOR dataset. This data contains genomic and clinical information obtained from 348 bladder cancer [MIM: 109800] patients ^34^. This analysis focuses on the pathway selection outcomes of GPS-Net while employing various reference pathway datasets. Three distinct reference pathway collections were employed: (1) hallmark gene sets collection (H), containing 50 gene pathways that emulate the conventional pathway selection problem with a small number of pathway; (2) cell type signature gene sets collection (C8), which contains 830 pathways, representing a larger-scale pathway referencing scenario; (3) Ecotyper cell state gene sets ^35^, consisting cell-type-specific reference signatures for 12 cell lineages, which have been shown associated with bladder cancer patients’ survival.

**Figure 4.**
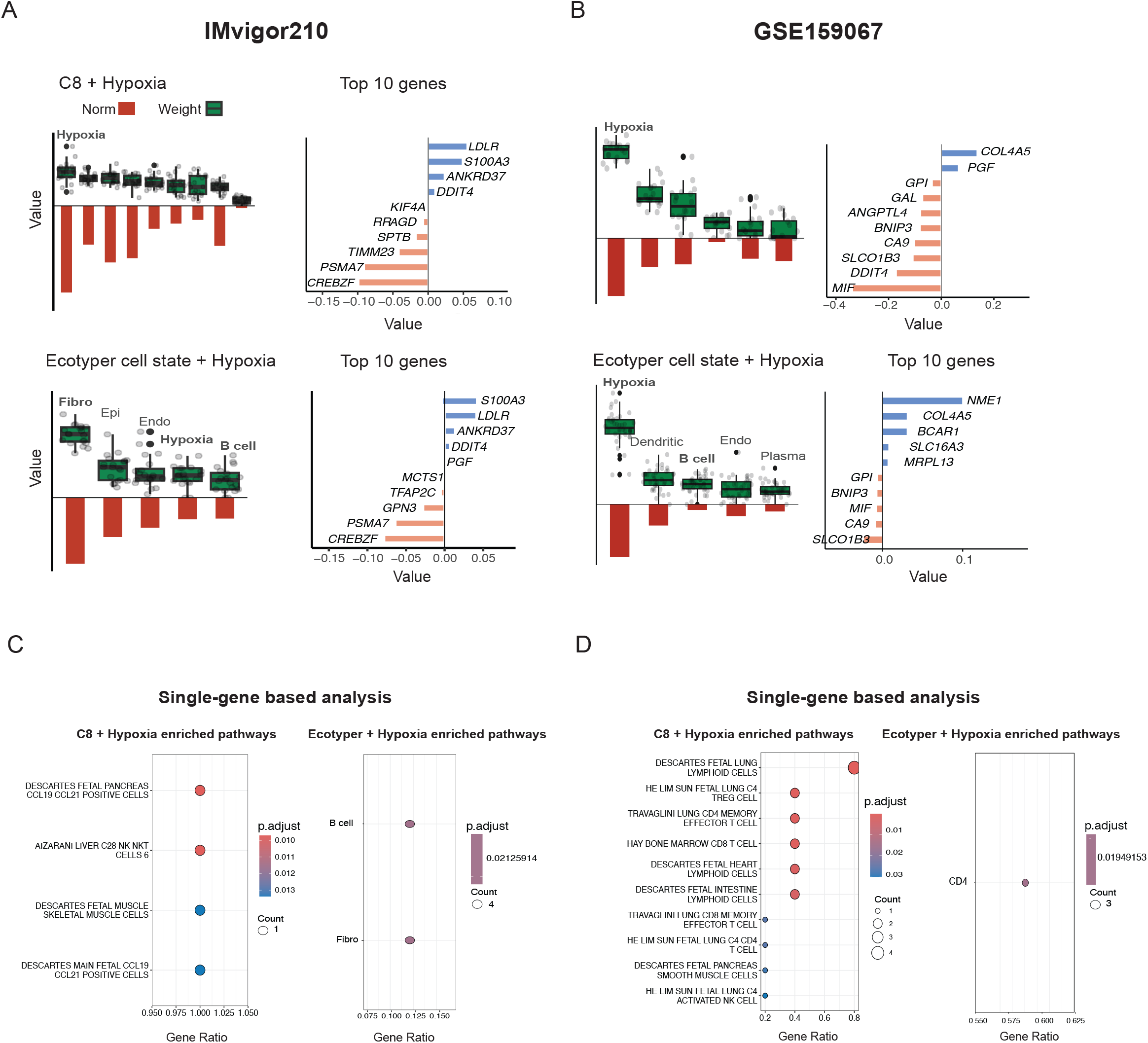
Prognostic pathway selection with immunotherapy datasets. Hypoxia pathway identification and top 10 genes selected in hypoxia pathway with different gene set backgrounds: (A) Bladder cancer from IMvigor201: C8 on the top and ecotype cell state on the bottom (B) HNSC from GSE159067: C8 on the top and ecotype cell state on the bottom. Pathway selection using single-gene based gene set enrichment method: (C) Bladder cancer (D) HNSC.

The estimated coefficients’ norms for pathways within the 50 hallmark gene sets and selected pathway names are shown in **Figure S2**. In this context, GPS-Net selected six pathways, with hallmark spermatogenesis emerging as the most significant pathway, marked by the highest norm value of 0.9. Notably, within this pathway, the norms of genes also display the largest values, reaching 0.4 in ANet’s estimations. However, when ANet selected genes within pathways also favored by GPS-Net, we observe a reduction in the norms of these coefficients. This is due to ANet’s broader selection of genes in pathways not chosen by GPS-Net. Three pathways were selected by both GPS-Net and Glasso, indicating a commonality between different selection methods. Yet, Glasso selected three additional pathways, two of which did not include any genes selected by Anet, highlighting a divergence in the selections made by different methods. A key distinction in GPS-Net’s approach was its consistent emphasis on the significance of entire pathways, a feature that gene-level section method ANet lacks. On the other hand, without incorporating the biological network information, Glasso tended to select more noisy pathways.

Next, we further assessed the pathway selection performance of GPS-Net using reference data. Specifically, we introduce the well-documented and significant hypoxia pathway, alongside the C8 and Ecotyper gene sets collection. In bladder cancer, hypoxia pathway’s activation can influence the behavior of cancer cells, contributing to their survival and aggressive features ^36,37^. The aim was to examine if GPS-Net could effectively identify the hypoxia pathway within the context of reference pathway data. As depicted in Figure 5B, we evaluated the pathway weights and the norms of gene coefficients within the selected pathways according to GPS-Net estimations. In the C8 reference background, the hypoxia pathway emerged as the most notable among the nine selected pathways. Additionally, the norms of genes within the hypoxia pathway shows the highest values. This demonstrates that GPS-Net effectively recognizes the importance of the hypoxia pathway, even when placed within a reasonably extensive collection of pathways (860 in total) as the reference background. In the selection with the Ecotyper pathway reference, the hypoxia pathway ranked fourth among the five selected pathways. The Ecotyper pathway collection includes genes that are already established as significantly associated with bladder cancer. This result underscores GPS-Net’s capacity to identify hypoxia even among pathways that are also evidently significant. Figure 4A further illustrates the top genes identified by GPS-Net in the hypoxia pathway with different reference gene sets C8 and Ecotyper gene sets, respectively. Notably, several genes exhibit consistent selection patterns in both reference cases. *LDLR* ([MIM: 606945]), *S100A3* ([MIM: 176992]), *ANKRD37* ([MIM: 619021]), and *DDIT4* ([MIM: 607729]), are among the genes consistently selected with positive coefficients, indicating a positive correlation with survival risk. On the contrary, *CREBZF* ([MIM: 606444]) and *PSMA7* ([MIM: 606607]) consistently exhibit the most negative association with survival risk in both C8 and Ecotyper gene set references. These findings demonstrate the robustness and reliability of GPS-Net in identifying key genes associated with survival outcomes across different reference backgrounds.

In addition, we applied GPS-Net to a Head and Neck Squamous Cell Carcinoma (HNSC) [MIM: 275355] gene expression dataset (GSE159067). This dataset contains 102 HNSC tumors and 2,559 genes. We used GPS-Net to select from the combination of hypoxia and C8 pathways, and hypoxia pathway emerged as the most significant as expected (Figure 4B). We further tested GPS-Net against the combination of hypoxia and ecotype cell state gene sets, and hypoxia remained the top selected pathway. The top 10 genes from the C8 background and ecotype cell state gene sets background showed consistent associations with survival, demonstrating the reliability of GPS-Net in both pathway and gene-level selection.

We further compared GPS-Net with a single-gene-based method that uses gene set enrichment analysis to identify significant pathways (Figures 4C and 4D). Both datasets revealed that the single-gene-based method failed to select the hypoxia pathway. Moreover, when searching for significant genes with a p-value less than 0.05, gene set enrichment analysis often failed to identify any pathways in high-dimensional gene expression data.

In summary, our study provides validation for GPS-Net’s pathway selection abilities within two distinct contexts. Using hallmark collection, we demonstrate that GPS-Net advances in identifying substantial and biologically relevant pathways. Using reference data, we further illustrate GPS-Net consistently recognize the critical hypoxia pathway, even within an extensive reference background. This suggests GPS-Net’s robustness in identifying key pathways among significant collections.

## Discussion

In this study, we propose the GPS-Net framework, an efficient tool for enhanced pathway selection, especially within survival data. The strength of GPS-Net lies in its innovative pathway kernel approach, a novel concept that seamlessly incorporates multiple pathway information into the selection process. This approach quantifies the pairwise interactions among genes within pathways by the application of Laplacian matrix, facilitating the integration of pathway-level insights into its selection strategy. In addition, the PK approach strategically tackles the computational burdens associated with the selection among a large number of pathways. This key property of GPS-Net significantly enhances our understanding of how these pathways and genes collaboratively function in genomic processes. We investigate capabilities of the GPS-Net in various simulation scenarios and read data applications.

Our study highlights GPS-Net’s comparative effectiveness when evaluated against current state-of-the-art biomarker selection techniques. Through a series of simulation scenarios, we have showcased its unique strength in genome-wide pathway selection. GPS-Net’s utilization of biological pathway network information consistently outperformed other methods in both marker and pathway selection. Additionally, our study illustrated GPS-Net’s proficiency in recognizing biologically significant pathway (hypoxia) within varying reference pathway backgrounds (H and C8). Furthermore, GPS-Net demonstrated its capability in quantifying the importance of significant pathways when substantial pathway reference is used (hypoxia + ecotype cell state pathways).

Constructing kernels from pathways in genomic data comes with its challenges, particularly the handling of gene overlaps between pathways. In our experiments, we reconstruct the pathways without overlap genes for fair comparison between GPS-Net and Glasso. While when there are gene overlaps between two pathways, the re-constructed kernels with overlap genes can be directly applied in GPS-Net framework. This is because the formulation does not require exclusive pathway representations, allowing pathways to coexist harmoniously without introducing significant redundancy. However, when gene overlaps exceed the 40% among the total number within pathways, it is often suggested to merge these pathways into a single new pathway ^38,39^. This consolidation helps reduce redundancy carried by multiple kernels, especially in managing complex genomic datasets. Researchers also have the option to re-construct pathways based on their insights and experiences, which can be a straightforward strategy for avoiding high overlap issues.

In GPS-Net, the utilization of graph kernels based on the Laplacian matrix is a prominent strategy for harnessing network information in pathway selection. When combined with the pathway kernel (PK) approach in GPS-Net, the MKL process aligns directly with the original task of pathway and biomarker selection. However, it is important to note that alternative kernels can be applied in this context. The choice of kernel has a substantial impact on the results and the performance of the final pathway selection. By considering various kernels, like polynomial and RBF kernels alongside graph kernels, a broader spectrum of pathway network structures can potentially be captured. Nevertheless, integrating different kernels requires tackling a new transformation from multiple pathway selection (4) to MKL problem, which is not as straightforward as using graph kernels. It also requires users to have a deeper understanding of kernel concepts. Another challenge in the application of MKL to pathway selection arises when dealing with large sample sizes. MKL excels in managing numerous pathways or genes (M and P), but it faces limitations in handling large sample sizes. A potential solution involves refining the GPS-Net algorithm by applying techniques like low-rank kernel learning ^40,41^, thus simplifying the computation when dealing with larger sample sizes. A recent study utilizing projected dual preconditioned SGD for large-scale general kernel models, in conjunction with Neural Tangent Kernels (NTK), provide new potential for the MKL model to efficiently handle large datasets by leveraging deep learning frameworks. ^33,42^

In our observations from simulation Scenario 2, it is evident that an accurate network structure significantly enhances pathway selection performance. Notably, recent studies focusing on integrating biological insights into high-dimensional data using integrative clustering ^43^ have encountered similar challenges in determining suitable network structures. For instance, the Generalized Bayesian Factor Analysis (GBFA) framework ^44^ has leveraged prior structural knowledge, such as the Laplacian prior, to facilitate feature selection guided by gene networks. GBFA transforms pathway information into a lower-dimensional latent factor loading space, thereby improving interpretability and relevance. To improve the estimation of the Laplacian matrix, the method used to derive the Laplacian matrix in GPS-Net can be applied in GBFA framework. This entails integrating data-driven and prior biological networks to obtain the final network representation.

Notably, the weights assigned to data-driven (reference-free) and prior biological (reference-based) networks can significantly impact pathway and gene set selection. In GPS-Net, users have the option to either fix this weight or determine it through cross-validation. The default setting for the weight is 0.5, reflecting the equal importance of network structures derived from data and prior knowledge. Assigning more weight to reference-free networks could lead to overfitting and a lack of biological interpretability. For different datasets, we recommend starting with the default weight as a benchmark, then refining the model through cross-validation.

In summary, the proposed GPS-Net framework improves network embedding, effectively addresses global gene-set selection challenges, and enhances the accuracy of prognostic biomarker and pathway identification. By leveraging both data-driven and existing gene-network information, GPS-Net demonstrates robust performance across various simulation scenarios and real-world datasets, making it a powerful tool for large-scale genomic research and precision medicine.

## Author contributions

Conceptualization, S.Y. and X.W.; Methodology, S.Y., K.L., P.F.K, and X.W; Investigation, S.Y., K.L., T. L., and X.W; Manuscript Writing, Review & Editing, S.Y., K.L., T. L., X.Y., P.F.K, and X.W.

## Acknowledgements

This work has been supported in part by a National Institutes of Health grant [R01DE030493 to X.W]; and Biostatistics and Bioinformatics Shared Resources at the H. Lee Moffitt Cancer Center *p* Research Institute, an NCI-designed Comprehensive Cancer Center (P30-CA076292).

## Data and code availability

GPS-Net is implemented in an R package and is available from https://github.com/wanglab1/GPSNet.

