## Supplementary Information for "GPS-Net: discovering prognostic pathway modules based on network regularized kernel learning"

- **Supplemental figures**
- **Supplemental tables**
- **Supplemental methods**

### Supplemental figures

**Figure S1**: Hypoxia selection with simulated pathways M=5 and M=10

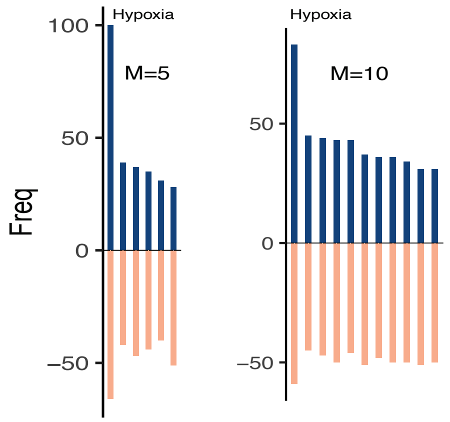

**Figure S2:** Hallmark selection in IMvigor210

A

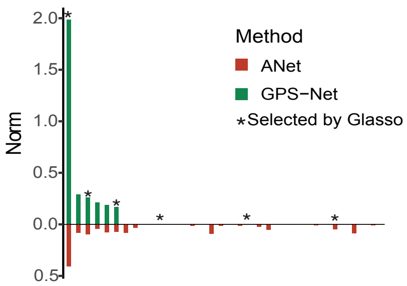


B

**Selected pathway names of GPS-Net for IMvigor210 data**

| Rank | Selected Hallmark pathway |
|---|---|
| 1 | HALLMARK_SPERMATOGENESIS |
| 2 | HALLMARK_E2F_TARGETS |
| 3 | HALLMARK_ESTROGEN_RESPNSE_EARLY |
| 4 | HALLMARK_TNFA_SIGNALING_VIA_NFKB |
| 5 | HALLMARK_INTERFERON_GAMMA_RESPONSE |
| 6 | HALLMARK_ESTROGEN_RESPONSE_LATE |

(A) Hallmarkpathway selection norms: GPS-Net's pathway selection results, displaying estimated norms for each pathway from the hallmark collection (50 pathways). Comparison with ANet's estimations and Glasso'spathway selections.
(B) Selected hallmark pathway names from GPS-Net.

### Supplemental tables

**Table S1**: Simulation settings with real pathway structure

| Pathway type | Pathway number | Simulation pathway number | Median gene number in each pathway | Simulation gene number | Simulation genes |
|---|---|---|---|---|---|
| H allmark | 50 | 50 | 180 | 200 | 10000 |
| C 1 | 301 | 500 | 115 | 100 | 50000 |
| C 8 | 829 | 1000 | 115 | 100 | 20000 |
| G o | 10460 | 10000 | 18 | 20 | 20000 |

**Table S2**: Scenario 5: Type I error with permutation test

| Generating model | Pathway setting | Significance level | GPS-Net | Anet | Enet | Lasso | Glasso |
|---|---|---|---|---|---|---|---|
| Null model: all $\beta = 0$ | 50 | 0.001 | 0.0014 | 0.0013 | 0.0021 | 0.0023 | 0.0024 |
| | | 0.01 | 0.0092 | 0.0086 | 0.0123 | 0.0130 | 0.0135 |
| | | 0.05 | 0.0422 | 0.0416 | 0.0485 | 0.0490 | 0.0489 |
| | 500 | 0.001 | 0.0012 | 0.0013 | 0.0021 | 0.0022 | 0.0023 |
| | | 0.01 | 0.0083 | 0.0079 | 0.0093 | 0.0094 | 0.0095 |
| | | 0.05 | 0.0404 | 0.0393 | 0.0474 | 0.0475 | 0.0476 |
| | 1000 | 0.001 | 0.0012 | 0.0014 | 0.0021 | 0.0022 | 0.0024 |
| | | 0.01 | 0.0084 | 0.0085 | 0.0095 | 0.0096 | 0.0096 |
| | | 0.05 | 0.0401 | 0.0354 | 0.0455 | 0.0456 | 0.0473 |
| | 10000 | 0.001 | 0.0013 | 0.0015 | 0.0020 | 0.0021 | 0.0023 |
| | | 0.01 | 0.0085 | 0.0090 | 0.0100 | 0.0101 | 0.0103 |
| | | 0.05 | 0.0405 | 0.0360 | 0.0460 | 0.0461 | 0.0485 |
| Mixed model: $\beta \neq 0$ for 5 pathways | 50 | 0.001 | 0.0023 | 0.0021 | 0.0043 | 0.0045 | 0.0056 |
| | | 0.01 | 0.0193 | 0.0184 | 0.0354 | 0.0360 | 0.0253 |
| | | 0.05 | 0.0554 | 0.0543 | 0.0665 | 0.0670 | 0.0724 |
| | 500 | 0.001 | 0.0153 | 0.0144 | 0.0254 | 0.0256 | 0.0183 |
| | | 0.01 | 0.0234 | 0.0224 | 0.0404 | 0.0410 | 0.0305 |
| | | 0.05 | 0.0615 | 0.0594 | 0.0713 | 0.0720 | 0.0805 |
| | 1000 | 0.001 | 0.0165 | 0.0184 | 0.0194 | 0.0196 | 0.0253 |
| | | 0.01 | 0.0263 | 0.0297 | 0.0324 | 0.0356 | 0.0405 |
| | | 0.05 | 0.0754 | 0.0758 | 0.0813 | 0.0820 | 0.0915 |
| | 10000 | 0.001 | 0.0184 | 0.0184 | 0.0195 | 0.0196 | 0.0335 |
| | | 0.01 | 0.0265 | 0.0295 | 0.0325 | 0.03527 | 0.0456 |
| | | 0.05 | 0.0755 | 0.0757 | 0.0820 | 0.0827 | 0.1156 |

### Supplemental methods

#### A.1 Normalized Laplacian matrix

The normalized Laplacian matrix represents the network connectivity and interactions between genes within a pathway. In normalized Laplacian matrix, the self-gene connection is 1 which corresponds to the value of $L_{ii} = 1$. The connection between gene i and gene j is described by the value of

$$L_{ij} = -\frac{c_{ij}}{\sqrt{\deg(i)\deg(j)}},$$

with $c_{ij}$ being the connectivity strength between gene i and gene j. $\deg(i)$ is the degree of the gene i which represents the number of genes connected to gene i. Mathematically, deg(i) is defined as the row sum of the corresponding adjacency matrix. The Laplacian matrix is defined as $L = D - A$, where $\mathrm{D}$ is the degree matrix and $\mathrm{A}$ is the adjacency matrix. If $A_{ij} = 1$, it indicates that there is an edge between gene iii and gene j. Therefore, the sum of the i-th row of $A$ corresponds to the number of genes connected to gene i by an edge. By definition, this sum is exactly the number of genes connected to gene i.

#### A.2 Solution for network regularization model (problem (4) and (5))

First, we introduce the procedure for the network regularization problem (4). For the binary response and survival outcome, a quadratic approximation of the log likelihood using secondary Taylor expansion at $a = \tilde{\beta}_0\mathbf{1} + X\tilde{\beta}$ ($\tilde{\beta}_0 = 0$ for Cox model) is applied:

$$\tilde{\ell}(\beta) = \frac{1}{2}(z - \beta_0\mathbf{1} - X\beta)^T \tilde{\ell}''_a (z - \beta_0\mathbf{1} - X\beta) + U(\tilde{\beta}),$$

where $z = a - \left(\tilde{\ell}''_a\right)^{-1}\tilde{\ell}'_a,\ U(\tilde{\beta})$ does not depend on $\beta$ ($\beta_0 = 0$ for Cox model). To speed up the computation, we assume the zero values for off-diagonal entries of $\tilde{\ell}''_a$ (Simon et al., 2011). For logistic model, $\tilde{\ell}''_a$ is a diagonal matrix. Let $u$ be the diagonal elements of $\tilde{\ell}''_a$,

$$\tilde{\ell}(\beta) \approx \frac{1}{2}\sum_{i=1}^{n} u_i (z - \beta_0\mathbf{1} - X\beta)^2.$$

The proximal Newton based coordinate ascent algorithm (Friedman et al., 2010; Simon et al., 2011) is applied,

**Pseudo code 1: for solving $\underset{\boldsymbol{\beta}}{\mathbf{argmin}}\left\{-\boldsymbol{\ell}(\boldsymbol{\beta}) + \boldsymbol{\lambda_1}\|\boldsymbol{\beta}\| + \frac{\lambda_2}{2}\boldsymbol{\beta}'\boldsymbol{L}\boldsymbol{\beta}\right\}$**

**Input:** gene expression data $\boldsymbol{X}$, response $\boldsymbol{Y}$, normalized Laplacian matrix $\boldsymbol{L}$, tolerance $\boldsymbol{\varepsilon}$, regularization parameters $\boldsymbol{\lambda_1}, \boldsymbol{\lambda_2} > \mathbf{0}$

**while** $||\boldsymbol{\beta}^{(t+1)} - \boldsymbol{\beta}^{(t)}|| > \boldsymbol{\varepsilon}$, **do**

$$\boldsymbol{\beta}_j^{(t+1)} \leftarrow \frac{S(\boldsymbol{A}^{(t)} - \boldsymbol{B}^{(t)}, \lambda_1 s_j)}{\frac{1}{n}\sum_{i=1}^{n} \boldsymbol{d}_i^{(t)} x_{ij} + \lambda_2 L_{jj}}$$

**end while**

$\boldsymbol{A}^{(t)} = \frac{1}{n}\sum_{i=1}^{n} \boldsymbol{u}_i^{(t)} \boldsymbol{x}_{ij}(\boldsymbol{z}_i^{(t)} - \boldsymbol{\beta}_0^{(t)} - \sum_{k\neq j} \boldsymbol{x}_{ik}\boldsymbol{\beta}_k^{(t)})$ , $\boldsymbol{B}^{(t)} = \boldsymbol{\lambda}_2 \sum_{k\neq j} \boldsymbol{L}_{jk}\boldsymbol{\beta}_k^{(t)}$ .

Function S(a,b) is the soft thresholding operator:

$$S(a, b) = sign(a)(|a| - b)_+ .$$

The different updating details is given by the following:
logistic model

$$\mu_i^{(t)} = \frac{1}{1+\exp(-X\beta^{(t)})}, z_i^{(t)} = \beta_0^{(t)} + x_i'\beta^{(t)} + \frac{y_i - \mu_i^{(t)}}{\mu_i^{(t)}(1-\mu_i^{(t)})},$$

$$u_i^{(t)} = \mu_i^{(t)}\left(1-\mu_i^{(t)}\right).$$

Cox model

$$a_i^{(t)} = X\beta^{(t)}, z_i^{(t)} = a_i^{(t)} + \frac{1}{u_i^{(t)}}\left[\delta_i - \sum \frac{d_i e^{a_i^{(t)}}}{\sum_{k\in R_j} e^{a_i^{(t)}}}\right],$$

$$u_i^{(t)} = \sum \frac{d_i e^{a_i^{(t)}}\left[\sum_{k\in R_j} e^{a_i^{(t)}} - e^{a_i^{(t)}}\right]}{\left(\sum_{k\in R_j} e^{a_i^{(t)}}\right)^2}.$$

Notice here, the algorithm can also handle the continuous responses with Gaussian model by $u_i = 1$ and $z_i = y_i$.

Second, we introduce the solution for mixture network regularization model problem (5). Consider a special case that $\lambda_1 = 0$ in (4). The Gaussian model for continuous responses in this case have the close form solution:

$$\hat{\beta} = (X'X + n\lambda_2 L)^{-1}X'Y.$$

We use kernel method in this case to speed up the computation for logistic and Cox model.

**Pseudo code 2: kernel method for (4) when $\lambda_1 = 0$**

1: Define the kernel matrix: $K = \frac{1}{n\lambda_2}XL^{-1}X'$.

2: Define the kernel transferred coefficient $\alpha$, $\alpha$ satisfies $\beta = \frac{1}{n\lambda_2}L^{-1}X'\alpha$.

3: The problem is transferred from $\hat{\beta} = \underset{\beta}{\operatorname{argmin}} -\frac{1}{2n}\ell(\beta) + \frac{\lambda_2}{2}\beta' L\beta$ to

$$\hat{\alpha} = \underset{\alpha}{\operatorname{argmin}} -\frac{1}{2n}\ell(\alpha) + \frac{\lambda_2}{2}\alpha' K\alpha$$

4: Solve the $\hat{\alpha}$ with the Newton update:

$$\alpha^{(t+1)} = \begin{cases} \left(U^{(t)}K + I\right)^{-1}\left(U^{(t)}K\alpha^{(t)} + Y - \mu^{(t)}\right), & \text{logisitc} \\ \left(V^{(t)}K + I\right)^{-1}\left(V^{(t)}L\alpha^{(t)} + W^{(t)}\right), & \text{Cox} \end{cases}$$

5: Final estimation: $\hat{\beta} = \frac{1}{n\lambda_2}L^{-1}X'\hat{\alpha}$.

For logistic model, $\mu^{(t)} = \frac{1}{[1+\exp(-K\alpha^{(t)})]}$ and $U^{(t)} = \text{diag}\{\mu^{(t)} * (1-\mu^{(t)})\}$.

For Cox model, $V^{(t)} = \text{diag}\{\exp(K\alpha^{(t)}) * H^{(t)}\}$, $H_i^{(t)} = \sum_{t_j \le t_i} h_j^{(t)}$, $h_j^{(t)} = \frac{1}{\sum_{j\in R_i}\exp(K\alpha^{(t)})}$ and $W^{(t)} = \delta - \exp(K\alpha^{(t)}) * H^{(t)}$. Here $*$ denotes the elementwise multiplication.

Solution for $(\hat{\beta}, \hat{w}) = \underset{\beta,w}{\operatorname{argmin}} \left\{-\ell(\beta) + \lambda_1\|\beta\| + \frac{\lambda_2}{2}\beta'[wL_1 + (1-w)L_2]\beta\right\}$:
Step1: Use pseudo code 2 to search for optimal $\hat{w} \in \{0,0.1,0.2,\dots,1\}$ with cross-validation.
Step2: Use pseudo code 1 to calculate the $\hat{\beta}_{\lambda_1\lambda_2}$ with L= $\hat{w}L_1 + (1-\hat{w})L_2$. Select best $\hat{\beta}_{\lambda_1\lambda_2}$ with cross-validation.

#### A.3 Dual problem and modified SpicyMKL with PK approach

In this section, we describe how PK approach is working for problem (8). The problem is solved by dealing with its dual problem. We begin with the introduction of convex conjugate function which is main concept of the dual problem. The convex conjugate function $c^*(\rho)$ of $c(x)$ is defined by,

$$c^*(\rho) = \sup_x[(\rho)'x - c(x)].$$

By Fenchel's duality theorem (Rockafellar 2015), if the regularity conditions are satisfied,

$$\inf_x[c_1(x) + c_2(x)] = \sup_y[-c_1^*(y) - c_2^*(y)], \qquad (A.3.1)$$

$c_1(\cdot)$ and $c_2(\cdot)$ are convex functions. Let $c_1(\alpha) = -\ell(\alpha)$ and $c_2(\alpha) = C\sum_{m=1}^{M}(1-\lambda)\|\alpha_m\|_{K_m} + \frac{\lambda}{2}\|\alpha_m\|_{K_m}^2 = \sum_{m=1}^{M}\phi_C(\alpha_m)$, by A.3.1 the problem (8) is then equivalent to the following,

$$\min_\alpha\left\{c_1(\alpha) + \sum_{m=1}^{M}\phi_C(\alpha_m)\right\} = \max_\rho\left\{-c_1^*(-\rho) - \sum_{m=1}^{M}\phi_c^*(\|\rho\|_{K_m})\right\}. \qquad (A.3.2)$$

The first order and second order partial derivatives of

$c_2(\alpha)$ is the summation of M convex functions and is still convex. The SpicyMKL algorithm [Suzuki et al., 2011] showed the gradient and hessian of $\phi_c^*(\|\rho\|_{K_m})$,

$$\nabla_\rho\left(\phi_c^*(\|\rho\|_{K_m})\right) = \frac{\phi_c^{(m)'}(\|\rho\|_{K_m})}{\|\rho\|_{K_m}}K_m\rho,$$

$$\nabla\nabla_\rho'\left(\phi_C{}^*\left(\|\rho\|_{K_m}\right)\right) = \phi_C{}'\left(\|\rho\|_{K_m}\right)\left(\frac{1}{\|\rho\|_{K_m}}K_m - \frac{1}{\|\rho\|_{K_m}^3}K_m\rho\rho^TK_m\right) + \frac{\phi_C{}''}{\|\rho\|_{K_m}^2}K_m\rho\rho^TK_m.$$

Where $\phi_c^{(m)'}(\|\rho\|_{K_m}) = \begin{cases} 0, & \|\rho\|_{K_m} \le C(1-\lambda) \\ \frac{\|\rho\|_{K_m} - C(1-\lambda)}{C\lambda}, & otherwise \end{cases}$ and $\phi_c^{(m)''}(\|\rho\|_{K_m}) = \begin{cases} 0, & \|\rho\|_{K_m} \le C(1-\lambda) \\ \frac{1}{C\lambda}, & otherwise \end{cases}$

Hence the gradient and hessian of A.3.2 are given by,

$$g_\rho = c_1^{*\prime}(-\rho) + \sum_{m=1}^{M}\nabla_\rho\left(\phi_c^*(\|\rho\|_{K_m})\right), \qquad (A.3.3)$$

$$H_\rho = c_1^{*\prime\prime}(-\rho) + \sum_{m=1}^{M}\nabla\nabla_\rho'\left(\phi_c^*(\|\rho\|_{K_m})\right). \qquad (A.3.4)$$

The algorithm for solving A.3.2 is given in the following,

**Pseudo code 3: Adjusted SpicyMKL for A.3.2**

1: Calculate $g_\rho$ and $H_\rho$ using A.3.3 and A.3.4.

2: Set the step size satisfying the constrain: $\eta^{(t)}$

3: Set the tolerance $\varepsilon$ .

4: Solve ρ with the Newton update:

while $||\rho^{(t+1)} - \rho^{(t)}|| > \varepsilon$, do

$\rho^{(t+1)} \leftarrow \rho^{(t)} + \eta^{(t)}\Delta;\ \ \Delta := -H_\rho{}^{-1} g_\rho$

5: $\hat{\alpha}_m = \frac{\phi c'\left(||\hat{\rho}||_{K_m}\right)}{||\hat{\rho}||_{K_m}} \hat{\rho}$.

6: $\hat{w}_m = \frac{\phi c'\left(||\hat{\rho}||_{K_m}\right)}{||\hat{\rho}||_{K_m}} / \sum \frac{\phi c'\left(||\hat{\rho}||_{K_j}\right)}{||\hat{\rho}||_{K_j}}$.

For logistic model, $\eta^{(t)}$ can be set following [Suzuki 2011]. And we showed the strategy to set the appropriate step size $\eta^{(t)}$ for Cox model in the following:

In Cox model, the following constrain is satisfied in each updating step,

$$\begin{cases} \rho_i < \delta_i, & 1 \le i \le n \\ \sum_{j=i}^{n} \rho_j < 0 & ,2 \le i \le n \end{cases}.$$

For each $\rho^{(t+1)}$, it should still satisfy this constrain. We define the following matrix A as,

$$A = \begin{pmatrix} 0 & 1 & 1 & \cdots & 1 \\ 0 & 0 & 1 & \cdots & 1 \\ \vdots & \vdots & \cdots & ⬚ & ⬚ \\ 0 & 0 & 0 & ⬚ & 1 \end{pmatrix},$$

$$A_{ij} = \begin{cases} 1, & j > i, or\ i = j = n \\ 0, & otherwise \end{cases}$$

Then the constrain becomes

$$\begin{cases} A\rho^{(t+1)} < 0 \\ \rho^{(t+1)} < \delta \end{cases} \Longrightarrow \begin{cases} A(\rho^{(t)} + \eta\Delta) < 0 \\ \rho^{(t)} + \eta\Delta < \delta \end{cases} \Longrightarrow \begin{cases} A\rho^{(t)} + \eta A\Delta < 0 \\ \rho^{(t)} + \eta\Delta < \delta \end{cases}$$

To find appropriate $\eta$, let $s_A = A\Delta$, we solved the following problem,

$$\begin{pmatrix} a_1 \\ \vdots \\ a_Q \end{pmatrix} + \eta \begin{pmatrix} b_1 \\ \vdots \\ b_Q \end{pmatrix} < \begin{pmatrix} c_1 \\ \vdots \\ c_Q \end{pmatrix},$$

With

$$\begin{pmatrix} a_1 \\ \vdots \\ a_Q \end{pmatrix} < \begin{pmatrix} c_1 \\ \vdots \\ c_Q \end{pmatrix}, and \begin{pmatrix} b_1 \\ \vdots \\ b_Q \end{pmatrix} > 0.$$

A solution can be found by $\eta = 0.9 \min_j \frac{c_j - a_j}{b_j}$. Using this method, we solve the $A\rho^{(t)} + \eta A\Delta < 0$ with $\eta_1$ and $\rho^{(t)} + \eta\Delta < \delta$ with $\eta_2$. The step size $\eta = \min\{\eta_1, \eta_2\}$.

We adjusted SpicyMKL by incorporating the Cox model to the original algorithm. We showed that the conjugate function of the Cox model is given by,

$$\ell^*(-\rho) = \begin{cases} \sum_{i=1}^{n}(\delta_i - \rho_i)\ln(\delta_i - \rho_i) - \sum_{i=1}^{n-1}\rho_i \ln \frac{\prod_{j=i+1}^{n}\left(-\sum_{m=j}^{n}\rho_m\right)}{\prod_{j=i+1}^{n} -\left(\sum_{m=j}^{n}\rho_m\right)}, & \rho_i < \delta_i, 1 \leq i \leq n \\ \sum_{i=1}^{n} - \delta_i \ln(\delta_i - \sum_{m=i}^{n}\rho_m) & , \sum_{m=i}^{n}\rho_m < 0, 2 \leq i \leq n \\ +\infty & , \quad else \end{cases}$$

And hence the gradient and hessian of $\ell^*(-\rho)$ is,

$$\ell_i^{*\prime} = \begin{cases} ln \frac{\prod_{j=i+1}^{n}(-\sum_{m=j}^{n}\rho_m)}{(\delta_i - \rho_i)\prod_{j=i+1}^{n}\delta_j - (\sum_{m=j}^{n}\rho_m)}, & i < n \\ -\ln(\delta_n - \rho_n) \quad , & i = n \end{cases}$$

$$\ell_{ij}^{*\prime\prime} = \begin{cases} \frac{1}{\delta_i - \rho_i}, & i = j \\ 0 \quad , & i \neq j \end{cases}.$$

**A.4 Adapted version of APM-$L_0$ in GPS-Net**

We augment a new parameter vector $\zeta_{\square}^{(\alpha)}$ which is defined by,

$$\zeta_m^{(\alpha)} = |\alpha_m|, m = 1, \ldots, M.$$

We add an extra $\lambda_3\left\|\zeta^{(\alpha)}\right\|_0$ penalty to (9) to eliminate the false positive groups in pathway selection. Since the $L_0$ norm does not have a dual form, we applied the APM-$L_0$ idea in the modified SpicyMKL updating strategies with convex approximation functions for $\lambda_3\left\|\zeta^{(\alpha)}\right\|_0$:

$$\min_{\alpha}\left\{c_1(\alpha) + \sum_{m=1}^{M}\phi_C\left(\alpha_m^{\square}\right) + \lambda_3\left\|\zeta^{(\alpha)}\right\|_0\right\} \; subject\ to \sum_{m=1}^{M}\left\|\alpha_m - \zeta_m^{(\alpha)}\right\|_2^2 \leq r$$

The updating algorithm is:

$$\alpha_m^{t+1} = \underset{\zeta_m^{(\alpha)}}{\operatorname{argmin}}\left\{c_1(\alpha) + \sum_{m=1}^{M}\phi_C\left(\alpha_m^{\square}\right) + \tau\sum_{m=1}^{M}\left\|\alpha_m^{\square} - \zeta_m^{(\alpha),t}\right\|_2^2\right\} \qquad (A.4.1)$$

$$\zeta_m^{(\alpha),t+1} = \underset{\zeta_m^{(\alpha)}}{\operatorname{argmin}}\left\{\lambda_3\left\|\zeta^{(\alpha)}\right\|_0 + \tau\sum_{m=1}^{M}\left\|\alpha_m^{t+1} - \zeta_m^{(\alpha)}\right\|_2^2\right\} \qquad (A.4.2)$$

Let $\phi_c^2\left(\alpha_m^{\square}\right) = \phi_C\left(\alpha_m^{\square}\right) + \tau\left\|\alpha_m^{\square} - \zeta_m^{(\alpha),t}\right\|_2^2$ and (A.4.1) can be solved by substituting $\phi_C\left(\alpha_m^{\square}\right)$ by $\phi_c^2\left(\alpha_m^{\square}\right)$ in $(A.3.2)$.

#### A.5 Estimating the reference-free and reference-based Laplacian matrices

**Pseudo code 4: estimating reference-free Laplacian matrix list in get_free_net**

```
1: Input: data matrix X (n by p); Output: normalized reference-free L
2: Load the ‘huge’ package in R
3: Estimate the precision matrix:
       fit<-huge(X, method=’glasso’)
       opt<-huge.select(fit)
       pre_mat<-fit$opt.icov
4: Convert the pre_mat to adjacency matrix A
5: Compute the degree matrix D with A
6: Compute the reference-free Laplacian matrix: L=D-A
7: Normalize L
```

**Pseudo code 5: estimating reference-based Laplacian matrix list in get_string_net**

```
1: Input: pathway gene list; Output: normalized reference-free L
2: Fetching interaction from STRING with R package “httr”:
     using httr::GET function from "https://string-db.org/api/tsv/network"
3: Calculate the adjacency matrix A from interaction data
4:   Convert the pre_mat to adjacency matrix
5: Compute the degree matrix D with A
6: Compute the reference-free Laplacian matrix: L=D-A
7: Normalize reference-based L
```

**A.6 Implementation workflow of (reference-based) GPS-Net**

**Pseudo code 6: Selection with provided Laplacian matrices (reference-free or reference-based)**

1: Input: data X, response Y, Laplacian matrices list L; Output: network weights w, estimator $\beta, \alpha, \rho$

2: Calculate Kernel matrices: $K_m = \frac{1}{M} X L_m^{-1} X'$

3: Apply **Pseudo code 3** to calculate MKL dual solution $\rho$

4: Compute MKL optimizer: $\hat{\alpha}_m = \frac{\phi_C'\left(||\hat{\rho}||_{K_m}\right)}{||\hat{\rho}||_{K_m}} \hat{\rho}$. Pathway selection weight: $\hat{w}_m = \frac{\phi_C'\left(||\hat{\rho}||_{K_m}\right)}{||\hat{\rho}||_{K_m}} / \sum \frac{\phi_C'\left(||\hat{\rho}||_{K_j}\right)}{||\hat{\rho}||_{K_j}}$.

5: Compute gene coefficient $\hat{\beta} = \sum_{m=1}^{M} L_m^{-1} X' \hat{\alpha}_m$

**Pseudo code 7: workflow of reference-based GPS-Net**

1: Input: data X, response Y, gene set (pathway) list; Output: network weights $\hat{w}_m^{(free)}$, $\hat{w}_m^{(ba)}$, estimator $\beta$

2: Get reference-free Laplacian matrix list with **Pseudo code 4**

3: Apply **Pseudo code 6** to calculate the integrated reference-free L: $\hat{L}^{(free)} = \sum_{m=1}^{M} \hat{w}_m^{(free)} L_m^{(free)}$

4: Apply **Pseudo code 5** to get reference-based Laplacian matrix list.

5: Apply **Pseudo code 6** to calculate the integrated reference-based L: $\hat{L}^{(ba)} = \sum_{m=1}^{M} \hat{w}_m^{(ba)} L_m^{(ba)}$

6: Apply **Pseudo code 1** to solve:

$$\underset{\beta}{\operatorname{argmin}} -\ell(\beta) + \lambda_1 \sum_{I=1}^{P} s_i\, |\beta_i| + \frac{\lambda_2}{2} \beta' \left[\gamma \hat{L}^{(free)} + (1-\gamma) \hat{L}^{(ba)}\right] \beta$$

$\gamma$: either use fixed value (default is 0.5) or tuning with cross-validation with **Pseudo code 2.**
